# Fast alignment and preprocessing of chromatin profiles with Chromap

**DOI:** 10.1101/2021.06.18.448995

**Authors:** Haowen Zhang, Li Song, Xiaotao Wang, Haoyu Cheng, Chenfei Wang, Clifford A. Meyer, Tao Liu, Ming Tang, Srinivas Aluru, Feng Yue, X. Shirley Liu, Heng Li

**Author notes:** These authors contributed equally to this work (Haowen Zhang, Li Song).

## Abstract

We present Chromap, an ultrafast method for aligning and preprocessing high throughput chromatin profiles. Chromap is comparable to BWA-MEM and Bowtie2 in alignment accuracy and is over 10 times faster than traditional workflows on bulk ChIP-seq / Hi-C profiles and than 10x Genomics’ CellRanger v2.0.0 pipeline on single-cell ATAC-seq profiles.

---

Chromatin profiling techniques, such as ChIP-seq^1^, ATAC-seq^2^, and Hi-C^3^, have been widely used to study transcription factor binding^4^, chromatin accessibility^5^, and higher order chromatin organization^6,7^, respectively. Single-cell ATAC-seq (scATAC-seq) further enables the profiling of cis-regulatory elements in individual cells^8^. Standard analysis workflows, such as those used by the ENCODE project^9^, start with read mapping by the popular short read aligner BWA-MEM^10^ or Bowtie2^11^, along with alignment sorting and deduplication by SAMtools^12^ and Picard^13^. These steps are the common bottlenecks which may take hours or days to complete, compared to the downstream analysis steps such as peak calling by MACS2^14^ which usually takes minutes. One reason for such inefficiency is that the comprehensive base-level alignment results for the purpose of variant calling are unnecessary for most chromatin biology studies. Furthermore, alignment filtering, deduplication, and other preprocessing steps are handled by different methods sequentially in a standard workflow, and each step requires parsing from compressed files. Such repeated I/O significantly increases the running time.

Our group previously developed minimap2^15^, an efficient read aligner based on the minimizer sketch. It was initially designed for long reads of high error rate and then extended for short accurate reads. Although a few times faster than FM-index-based short-read aligners such as BWA-MEM and Bowtie2, minimap2 more frequently misses short alignments that lack sufficient minimizer seeds. This becomes a severe issue in mapping scATAC-seq data when a large portion of the read sequence is used for barcoding and indexing, and the remaining genomic sequence in a read can be as short as 50bp. Moreover, minimap2 has to slowly scrutinize the alignment to resolve the high sequencing error rate inherent in the long reads, which could be unnecessary for the highly accurate Illumina short-read sequencing data.

In this study, we present an efficient read alignment and preprocessing method, named Chromap, based on the minimizer sketch (Fig. 1a). Chromap features a fast sorting-based procedure to generate mapping candidates and a fast alignment algorithm to pick the best candidate. To better handle short reads, Chromap considers every minimizer hit and uses the mate-pair information to rescue remaining missing alignments caused by the lack of low-frequency minimizers. Taking advantage of the observation that chromatin profiles are enriched only in a subset of the whole genome, Chromap caches the candidate read alignment locations in those regions to accelerate alignment of future reads containing the same minimizers. Besides read mapping, Chromap also incorporates sequencing adapter trimming, duplicate removal and scATAC-seq barcode correction, which further improves the processing efficiency (Online Methods). Chromap significantly reduces the computational time without losing accuracy.

**Figure 1.**
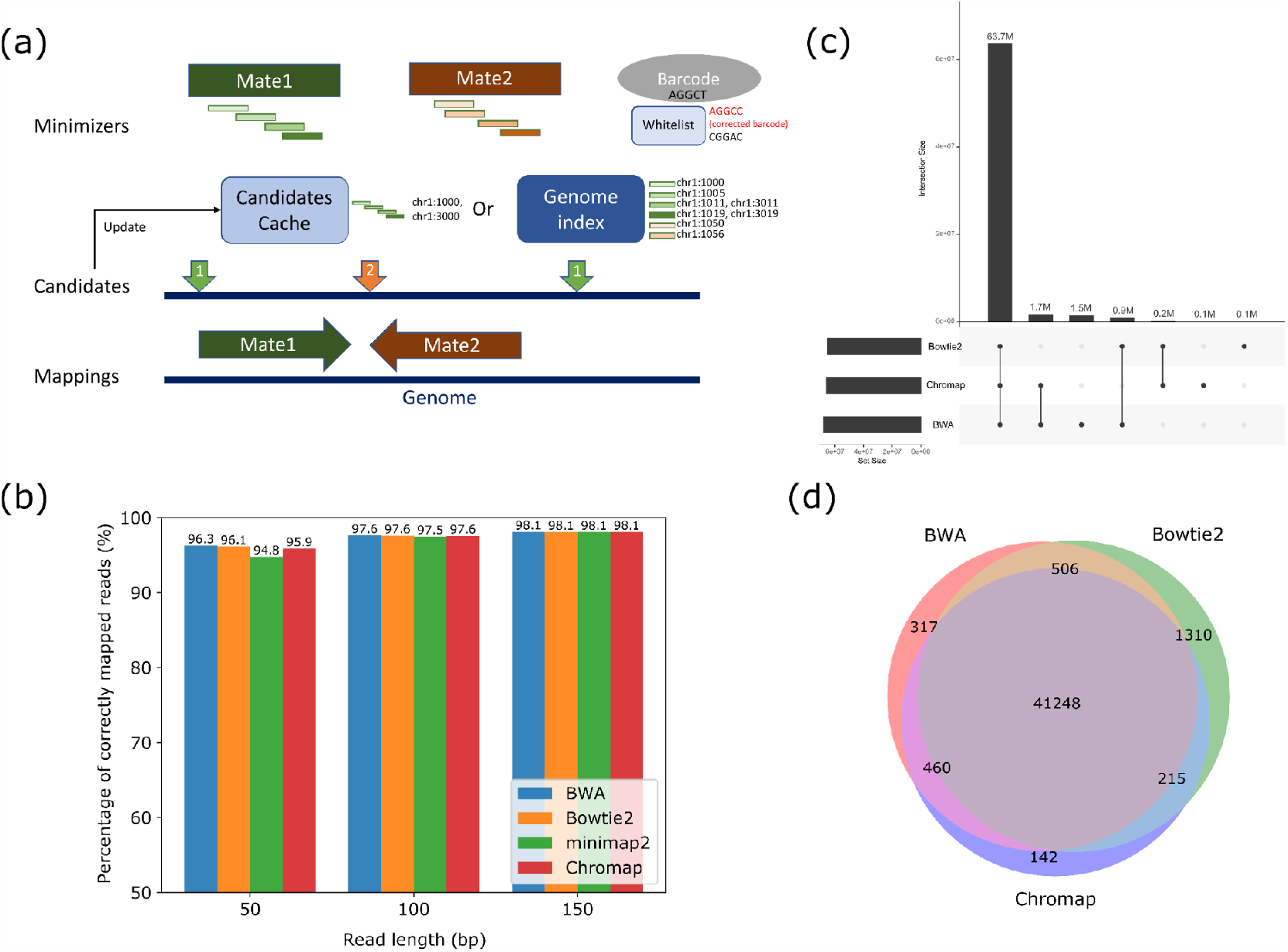
Overview of Chromap. (a) Workflow of Chromap (b) Accuracy of methods on the simulated data with different read length (c) Consensus of read alignments from Chromap, BWA-MEM and Bowtie2 on bulk ChIP-seq data (d) Overlapped peaks from the alignments file reported by different methods on bulk ChIP-seq data

We compared Chromap with BWA-MEM, Bowtie2, and minimap2 on three simulated whole genome sequencing data sets with various read lengths (Fig. 1b). The accuracy of these aligners was similar on the 100bp and 150bp paired-end data, about 98% for all three methods. On 50bp paired-end data, BWA-MEM, Bowtie2 and Chromap had similar accuracy of around 96%, while minimap2 had worse performance at 95%. The comparison confirmed the expected lower accuracy of minimap2 as it was not designed to align short reads. We thus excluded minimap2 from further benchmarking.

Next, we evaluated Chromap on a CTCF ChIP-seq data set from the ENCODE project and compared it with BWA-MEM and Bowtie2. Among the 68 million fragments reported by any of the three methods (MAPQ≥30), Chromap aligned 3% fewer fragments than BWA-MEM and 1.2% more than Bowtie2, and 99.8% of Chromap alignments were supported by either BWA-MEM or Bowtie2 (Fig. 1c). We next investigated the effects of the three alignment methods on peaks called by MACS2. Peaks from Chromap alignment overlapped 99.7% with those from BWA-MEM and Bowtie2, and Chromap created the fewest aligner-unique peaks (Fig. 1d). Annotation of the peaks with ChIPseeker^16^ did not find any aligner-specific bias in the three sets of peaks (Fig. S1). Overall, the differences from the three alignment methods were significantly smaller than those between data replicates (Fig. S2). Chromap only took less than 5 minutes to complete the mapping, sorting, and deduplication process, while the workflow based on BWA-MEM (or Bowtie2) and Picard required over 100 minutes. On the alignment step, Chromap (3.5min) was 18 times and 24 times faster than BWA-MEM (64min) and Bowtie2 (86min) respectively, supporting the efficiency improvement of Chromap. We note that Chromap also reduced half an hour on the sorting and deduplication steps, confirming the advantage of integrating alignment and preprocessing in chromatin profiling analysis.

Chromap supports split-alignment, thus is compatible with Hi-C analysis. We compared the performance of Chromap and BWA-MEM on a Hi-C data set on the K562^7^ cell line by evaluating the downstream chromatin features such as chromatin compartments, topologically associating domains (TADs), and chromatin loops. The chromatin compartments (measured by the first eigenvector) and TADs (measured by insulation score) called from the two aligners gave highly similar results, achieving Pearson correlation coefficients of 0.995 and 0.998 respectively (Fig. 2a, Fig. S3a). Although there is some divergence on the chromatin loops called by the two aligners, CTCF enrichment at the loop anchors supported these aligner-unique loops as genuine chromatin interaction loops (Fig. S5b, c. Online Methods). On this large data set with about 1.4 billion read pairs, Chromap spent 164 minutes to produce a processed alignment file in the pairs format^17^ ready for downstream analysis. It was 13 times faster than a standard workflow with BWA-MEM and pairtools^17^.

**Figure 2.**
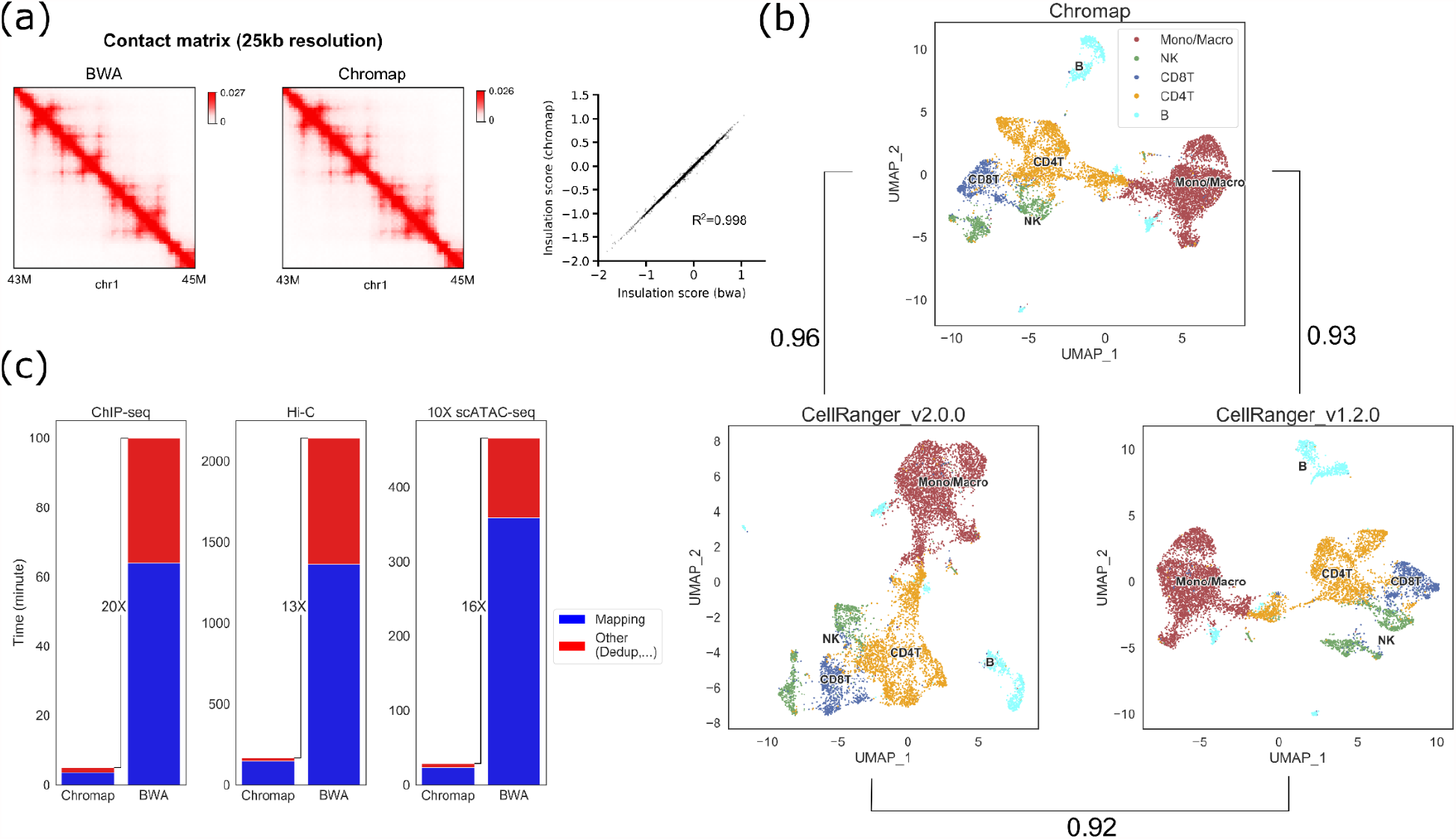
Chromap on the large data set. (a) Comparison of Hi-C contact matrices at 25kb resolution and insulation scores for TADs analysis derived from Chromap and BWA-MEM alignments. (b) Cluster annotation and NMI of the PBMC 10x Genomics scATAC-seq data based on the results of Chromap and CellRanger (c) Running time of Chromap and workflows based on BWA-MEM on ChIP-seq, Hi-C data and 10x Genomics scATAC-seq

Last but not least, we tested Chromap on a 10K PBMC scATAC-seq dataset from 10x Genomics with 750 million reads and compared the results with CellRanger v1.2.0 and CellRanger v2.0.0, the official pipelines for processing scATAC-seq data developed by 10x Genomics based on BWA-MEM. Released in May 2021, CellRanger v2.0.0 substantially improves the computational efficiency over its predecessor along with other updates in preprocessing steps, such as deduplication criteria. We used all three methods for alignment and preprocessing followed by MAESTRO^18^ for cell clustering and cell type annotation (Fig. 2b). We evaluated the consistency of cell type annotation using normalized mutual information (NMI), and found Chromap and CellRanger v2.0.0 generated nearly identical results with NMI more than 0.96, higher than the NMI between the two CellRanger versions (Fig. 2b, Table S1, Online Methods). The lower consistency between CellRanger v1.2.0 and v2.0.0 suggested that alternating BWA-MEM and Chromap had less impact on the analysis than changing other preprocessing strategies. The clustering profiles were also highly similar between Chromap and CellRanger v2.0.0, no matter whether clustering was performed using the peak-based approach in MAESTRO or the bin-based approach in ArchR^19^ (Table S2). On performance, Chromap generated the final alignment file in less than 30 minutes. It was 68 times faster than CellRanger v1.2.0 (33 hours) and 16 times faster than CellRanger v2.0.0 (8 hours). The memory usage of Chromap is around 20GB. Since the memory usage is dependent on the index file size, it is stable with respect to sequencing depth and regardless of applications to ChIP-seq, Hi-C, or scATAC-seq.

In summary, Chromap implements an efficient and accurate alignment and processing method for chromatin profiles. It is significantly faster than general-purpose aligners by taking full advantage of the nature of chromatin studies, i.e. sequencing reads are enriched in peaks and read coordinate locations are more important for downstream analyses (Fig. 2c). Chromap further improves efficiency by integrating the adapter trimming, alignment deduplication, and barcode correction processing steps in the standard chromatin biology data workflows. With the decreasing cost of high throughput sequencing and increasing deeper sequencing coverage of chromatin profiles, Chromap will continue to expedite biological findings from chromatin studies in the future.

## Methods

Methods, evaluation details and data availability are available in the online methods section.

## Acknowledgments

We acknowledge the following funding sources for supporting this work: Breast Cancer Foundation [BCRF-20-100 to XSL] and National Institute of Health [R01HG010040, R01HG011139, and U01CA226196 to HL].

## Author contributions

H.Z., L.S., X.S.L. and H.L. conceived the project. H.Z., L.S., X.W., H.C. and H.L. designed the method. H.Z. and L.S. implemented the method. H.Z., L.S., X.W., H.C., C.W., C.M., X.S.L. and H.L. evaluated the method. H.Z., L.S., X.W., S.A., F.Y., X.S.L. and H.L. wrote the manuscript. All authors read and approved the final manuscript.

## Online methods

### Index construction

Double-strand minimizers^15^ of reference genomes are collected and indexed using a hash table with minimizer sequences as keys and their sorted order of occurrences along the reference as values (Fig. 1a). Due to the repetitive regions in the reference, some minimizers have high frequency, which can cause false positive mappings and reduce mapping speed significantly. Thus by default, we mask minimizers occurring >500 times on the reference.

### Adapter removal

For ATAC-seq or scATAC-seq, when a read contains the adapter sequence at the 3’-end, its fragment length can be shorter than the read length. To remove the adapters, for a pair of reads, if a prefix of one read has ≤1 Hamming distance compared with a suffix of the other read in the pair and the overlapped region is longer than a threshold *l*_*ovp*_, we trim the bases outside the overlap. We extract *l*_*ovp*_/2 long seeds from one read, find the hits of the seeds in the other reads and verify those hits. This algorithm accelerates the trimming step and still guarantees finding overlaps within Hamming distance of 1.

### Candidate generation

We define candidates for a read to be possible mapping start locations on the reference genome, which are estimated by exact minimizer hits (i.e., anchors) between the read and the reference. Formally, an anchor is a pair (*x, y*) where *x* denotes the minimizer start position on the reference and *y* denotes the minimizer start position on the read. Then the candidate can be estimated by this anchor as *x - y* . Co-linear anchors (i.e., chains) are a set of anchors that appear in ascending order in both the read and reference, which can be found by a dynamic programming algorithm^15^ in quadratic time with respect to the number of anchors. While this algorithm can robustly identify chains for noisy long reads (>1000bp with 5%∼10% error rate), we present a more efficient algorithm that can generate candidates for short reads with a low error rate. We generate candidates using all the anchors and then sort the candidates. During a linear scan on the sorted candidates, we merge the same candidates or candidates that have smaller than error threshold difference generated from multiple anchors. During the merging, Chromap records the multiplicity for each candidate, which is also the number of supporting anchors, and filters the candidates with fewer support than the user-defined threshold. For paired-end reads, chains were first generated for each end and then filtered by the fragment length constraint.

### Candidate cache

Chromap stores the raw candidates in a cache for frequent reads to avoid repeated candidate generation for reads from peak regions. The cache is a hash table, where the key is a vector of minimizers and the value is the candidates generated from the set of minimizers. The minimizers vector stores the *M*minimizers sequences *m*_*i*_ and the *M* - 1 offsets between adjacent minimizers *m*_*i*_. Chromap uses the function 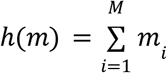 to map the vector to the *h(m)* entry in the hash table. The advantage of this mapping function is that the identical reads from both strands can access the same cached information. Furthermore, reads that are nearby in the genome have a greater likelihood of generating the same minimizer vector, and they can also share the same cache information.

Chromap maintains a small count array in each cache entry to identify the most frequent minimizer vector from hundreds of different vectors mapped to the same cache entry. Chromap uses the function *f(m) =* ⊕_*i*_*m*_*i*_ to map the vector to the *f(m)* entry in the count table by computing xor of the minimizer codings, which has the same advantage of ignoring the read strand. Chromap then updates the cache table if and only if the count for the minimizer vector is more than 20% of the total count in the count array, and is the dominant minimizer vector (show up more than half times) among the vectors mapped to the count array entry *f(m)*. As a result, Chromap not only stores in cache the candidates from frequent minimizers vectors, but also avoids unnecessary cache updates from the background noises.

### Candidate supplements

Chromap supplements the candidates with mate-pair information to recover the lost candidates due to the minimizer occurrence limit. For each read end, Chromap will pick the mate’s candidate supported by the most number of anchors and use this mate’s candidate as the estimation for the read coordinate. As a result, for each minimizer in the read end, instead of extracting all the occurrences on the reference, Chromap applies a binary search in the index entry to only select the occurrences within the range estimated read coordinate determined by the fragment size distribution. Chromap then executes the same candidate generation algorithm to supplement the candidates with the minimizer occurrences from the binary searches.

### Candidate verification

Since each read can have multiple candidate mapping positions, we implemented a banded Myers’ bit-parallel algorithm^20^ to pick the optimal candidate coordinate with minimum edit distance to the reference genome. To further accelerate the verification step, we parallelized the algorithm using SIMD instructions on the CPU to align the read with multiple candidate positions on the reference simultaneously. We also modified the algorithm to efficiently trace back the alignment so that accurate start and end mapping positions can be obtained.

### Split mapping

When the edit distance exceeds the threshold during the candidate verification step, we check if the length of the mapped read is greater than a certain length threshold. If the length of the mapping passes the length filter, the mapping is kept with an estimated mapping score as the mapped read length minus the edit distance. Note that for some of the Hi-C reads, there can be a small region (<20 bp) which cannot be mapped at the beginning of its 5’ end. To resolve this issue, when the mapping length is too short, the first 20 bp of the read is excluded and a second round of mapping of the remaining region is performed. If a mapping generated in this way passes the length filter, the mapping is then extended backward from its beginning to its maximum exact match. For paired-end data in split-alignment mode, Chromap ignores the constraints from the mate-pair, such as the fragment length or strandness.

### Deduplication

When the data set is small, all the mappings can be kept in the memory and sorted to remove duplicates. For large data sets or limited memory, we provide a low memory mode. It saves mappings in chunks temporarily on the disk and uses external sort to merge them into the final mapping output in a low memory footprint. For scATAC-seq data, duplicates can be removed at either bulk level or cell level (default) based on the users’ choice.

### Barcode correction

Using the barcode whitelist provided by 10x Genomics, we correct barcodes that are not on the whitelist. Prior to the correction, the barcodes are converted to their bit representations and the abundance of each barcode is computed efficiently using a hash table. For barcodes outside the whitelist, all whitelisted barcodes within one Hamming distance from the barcode to correct are extracted by a set of efficient bit operations. Using the quality score of the mismatched base and the abundance of these whitelisted barcodes as a priori, we compute the posterior probability of correcting the observed barcode to the whitelisted barcodes. We make the correction if the highest probability of the observed barcode being a real barcode is >= 90%. The correction step is performed during the mapping process and thus overlapped by read loading.

### Simulated and sequencing data for evaluation

In this work, we evaluated Chromap on various data sets including simulated whole genome sequencing data, bulk ChIP-seq data, 10x Genomics scATAC-seq data, and Hi-C data. One million fragments were simulated from the human reference genome GRCh38 using Mason^21^ with average sequencing error rate 0.1% and read lengths 50bp, 100bp, and 150bp. The bulk CTCF ChIP-seq data on the human VCaP cell line were downloaded from ENCODE to test the tools on bulk sequencing data. The 10K PBMC scATAC-seq dataset is publicly available from 10x Genomics and used to evaluate the performance of the tools on single cell data. To investigate the impact of alternating BWA-MEM with Chromap on chromatin conformation analysis, we combined the two Hi-C data replicates from a previous study^7^.

### Evaluating performance for ChIP-seq data

In this work, we mainly compared Chromap with two popular short read aligners BWA-MEM (v0.7.17) and Bowtie2 (v2.4.2). When testing on simulated data, we also included minimap2 (v2.17) in the evaluation and we converted all the alignments into PAF format and used the paftools to calculate the accuracy of alignments. Using the bulk ChIP-seq data, we compared the consensus of alignments and peaks among the three aligners after filtering the alignments with MAPQ less than 30 based on ENCODE protocol.

When evaluating the methods on Hi-C data, we filtered the alignments with MAPQ 0, which follows the default parameter settings in pairtools. All the methods were tested in a multiprocessing environment with 8 threads. CellRanger is a pipeline including data analysis steps after alignment and preprocessing, we measured its running time until the last “WRITE_ATAC_BAM” step in the log file.

### Evaluating performance for Hi-C data

We compared Chromap and the standard 4D Nucleome Hi-C processing pipeline, which is based on BWA-MEM and pairtools^17^, on a large Hi-C data set on human cell line K562. Due to complexity introduced by the ligation junction in a Hi-C experiment, direct comparison of alignment coordinates would underestimate the consistency between the methods. Therefore, we compared the contact maps derived from the alignments at various resolutions. We compared the overall distribution of chromatin contacts at 25kb resolution by using the stratum-adjusted correlation coefficients (SCC); the chromatin compartments measured by the first eigenvector of the normalized contact matrices at 100kb; the TAD boundary strength measured by the insulation score at 25kb resolution; and the identified chromatin loops at 10kb.

To confirm that the difference between Chromap alignments and BWA-MEM alignments was smaller than the difference between biological replicates, we computed SCCs by using a Python implementation of HiCRep (https://pypi.org/project/hicreppy/, v0.0.6)^22^ between Chromap and BWA-MEM on the same replicate (Chromap R2 vs. BWA-MEM R2) or between two replicates (BWA-MEM R1 vs. BWA-MEM R2). Because the original two replicates have disparate sequencing depths (R1: 1,048,612,352 vs. R2: 317,616,493), we first down-sampled R1 to make it match the sequencing depth of R2. The resulting SCC between Chromap R2 and BWA-MEM R2 was 0.998, which was significantly higher than SCC between BWA-MEM R1 and BWA-MEM R2 (0.945). HiCRep was run at 25kb resolution, and the smoothing factor and the maximum genomic distance were set to 5 and 2Mb, respectively. For the following chromatin conformation analysis, we merged the alignment results from the two replicates.

Both compartments and TADs were estimated using cooltools (https://pypi.org/project/cooltools/, v0.3.2). For compartments, the eigenvalue decomposition was performed on the 100kb intra-chromosomal contact maps and the first eigenvector (PC1) was used to capture the “plaid” contact pattern. The original PC1 was oriented according to a K562 DNase-Seq track (ENCODE accession code: ENCFF338LXW) so that positive values correspond to active genomic regions and negative values correspond to inactive regions. The Pearson correlation of PC1 was 0.995 between Chromap and BWA (Fig. S3a). For TADs, genome-wide insulation scores (IS) were calculated at 25kb with the window size setting to 1Mb. The Pearson correlation of the IS scores were 0.998 between the results from Chromap and BWA-MEM (Fig. 2b). Finally, we identified chromatin loops using HiCCUPS at 10kb (https://pypi.org/project/hicpeaks/, v0.3.4). Among the 9,455 and 9,950 loops identified from Chromap and BWA-MEM respectively, we found 8,385 of them were supported by both methods (Fig. S3b). Furthermore, we found loop anchor sites that were uniquely identified by Chromap or BWA-MEM had a similar enrichment of CTCF binding peaks (Fig. S3c), suggesting those aligner-specific anchors could be biologically meaningful.

### Evaluating performance for scATAC-seq data

We conducted comprehensive evaluations between Chromap and CellRanger on the 10K PBMC 10x Genomics scATAC-seq data to show that the clustering results were not affected by replacing CellRanger with Chromap. We compared the consistency of the cell type annotations or cell clusters using normalized mutual information (NMI) and adjusted rand index (ARI) calculated by the Python package scikit-learn. We first computed the baseline NMI and ARI between CellRanger v1.2.0 and CellRanger v2.0.0. Chromap vs CellRanger v2.0.0 achieved a higher consistency score than the baseline score, suggesting the results from Chromap were highly consistent with CellRanger and were more consistent than CellRanger version changes (Table S1). To confirm the difference in the consistency score was insignificant, we created two replicates of the data set by randomly sampling 95% of the read pairs in the data set and applied CellRanger v2.0.0 to process these two replicates. The cluster-level NMI between the two downsampled replicates (0.888) was lower to the NMI of the clusters generated from CellRanger v2.0.0 and Chromap (0.932), supporting that the impact from alternating CellRanger and Chromap is small. In addition, we also applied a bin-based scATAC-seq analysis method ArchR on this data set to evaluate the difference in the clustering caused by using Chromap and two CellRanger versions. Similar to the results on MAESTRO, we found alternating CellRanger to Chromap had tiny effects on the clustering results generated by ArchR (Table S2). Though CellRanger v1.2.0 is slow, it is easier to modify and we were able to adapt it to use Bowtie2 as the alignment method (CellRanger v1.2.0_Bt2). Therefore, we could examine the impact of alternating the alignment methods on cell clusters. In this case, we ran Chromap with bulk level deduplication (Chromap_bulkdedup) as the setting in CellRanger v1.2.0. The NMI and ARI scores among CellRanger v1.2.0, CellRanger v1.2.0_Bt2 and Chromap_bulkdedup are all high (NMI > 0.9, ARI>0.88, Table S3), suggesting that alternating the alignment methods BWA-MEM, Bowtie2 and Chromap had little impact on scATAC-seq analysis.

## Data availability

ChIP-seq data with two replicates are available from ENCODE: ENCSR265ARE 10x Genomics data is available at: https://support.10xgenomics.com/single-cell-atac/datasets/1.2.0/atac_v1_pbmc_10k Hi-C data is available in the SRA repositories SRR1658693∼SRR1658702.

The code used in the evaluation is available at: https://github.com/haowenz/chromap_evaluation Chromap source code is available at https://github.com/haowenz/chromap.

---

**Supplementary Table 1.** The normalized mutual information (NMI) and adjusted rand index (ARI) of cell type annotations and cell clusters from MAESTRO on 10K PBMC 10x Genomics scATAC-seq data using Chromap and CellRanger v1.2.0 and v2.0.0. MAESTRO obtained 15, 16, 15 clusters from CellRanger v1.2.0, CellRanger v2.0.0 and Chromap results respectively.

| Cluster |  |  |  |
| --- | --- | --- | --- |
|  | CellRanger_v1.2.0 vs<br>CellRanger_v2.0.0 | CellRanger_v1.2.0 vs<br>Chromap | CellRanger_v2.0.0 vs<br>Chromap |
| NMI | 0.822 | 0.832 | 0.932 |
| ARI | 0.677 | 0.701 | 0.914 |
| Cell type |  |  |  |
|  | CellRanger_v1.2.0 vs<br>CellRanger_v2.0.0 | CellRanger_v1.2.0 vs<br>Chromap | CellRanger_v2.0.0 vs<br>Chromap |
| NMI | 0.922 | 0.933 | 0.964 |
| ARI | 0.963 | 0.969 | 0.983 |

**Supplementary Table 2.** The normalized mutual information (NMI) and adjusted rand index (ARI) of cell type annotations and cell clusters from ArchR on 10K PBMC 10x Genomics scATAC-seq data. ArchR obtained 11, 12, 13 clusters from CellRanger v1.2.0, CellRanger v2.0.0 and Chromap results respectively.

| Cluster |  |  |  |
| --- | --- | --- | --- |
|  | CellRanger_v1.2.0 vs<br>CellRanger_v2.0.0 | CellRanger_v1.2.0<br>vs Chromap | CellRanger_v2.0.0<br>vs Chromap |
| NMI | 0.865 | 0.881 | 0.899 |
| ARI | 0.896 | 0.905 | 0.928 |

**Supplementary Table 3.**
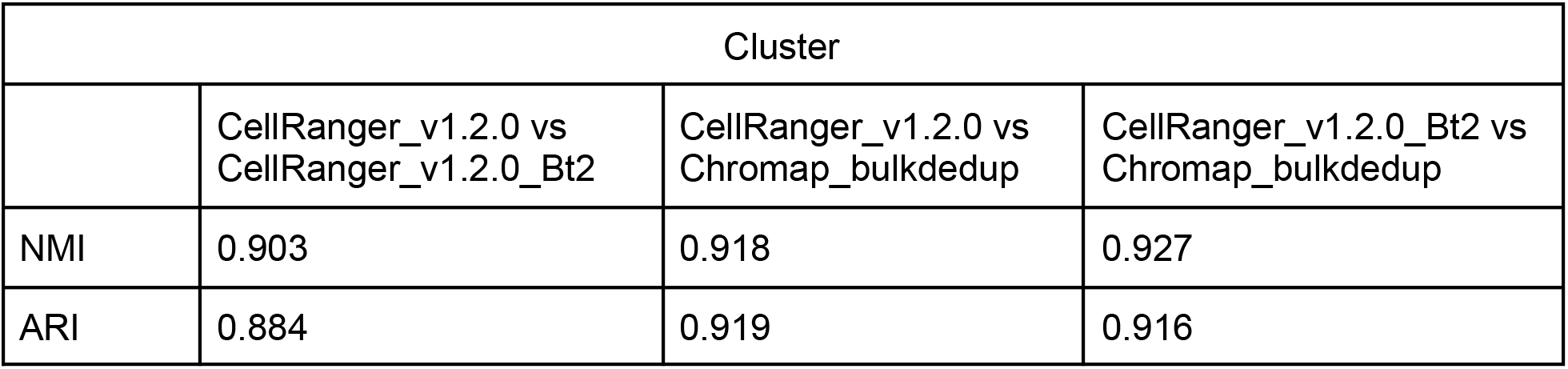
The normalized mutual information (NMI) and adjusted rand index (ARI) of cell clusters from MAESTRO on 10K PBMC 10x Genomics scATAC-seq data using Chromap_bulkdedup and CellRanger v1.2.0 with BWA and Bowtie2 as aligners MAESTRO obtained 15, 14, 15 clusters from BWA, Bowtie2 and Chromap results respectively.

| Cluster |  |  |  |
| --- | --- | --- | --- |
|  | CellRanger_v1.2.0 vs<br>CellRanger_v1.2.0_Bt2 | CellRanger_v1.2.0 vs<br>Chromap_bulkdedup | CellRanger_v1.2.0_Bt2 vs<br>Chromap_bulkdedup |
| NMI | 0.903 | 0.918 | 0.927 |
| ARI | 0.884 | 0.919 | 0.916 |

**Supplementary Figure 1.**
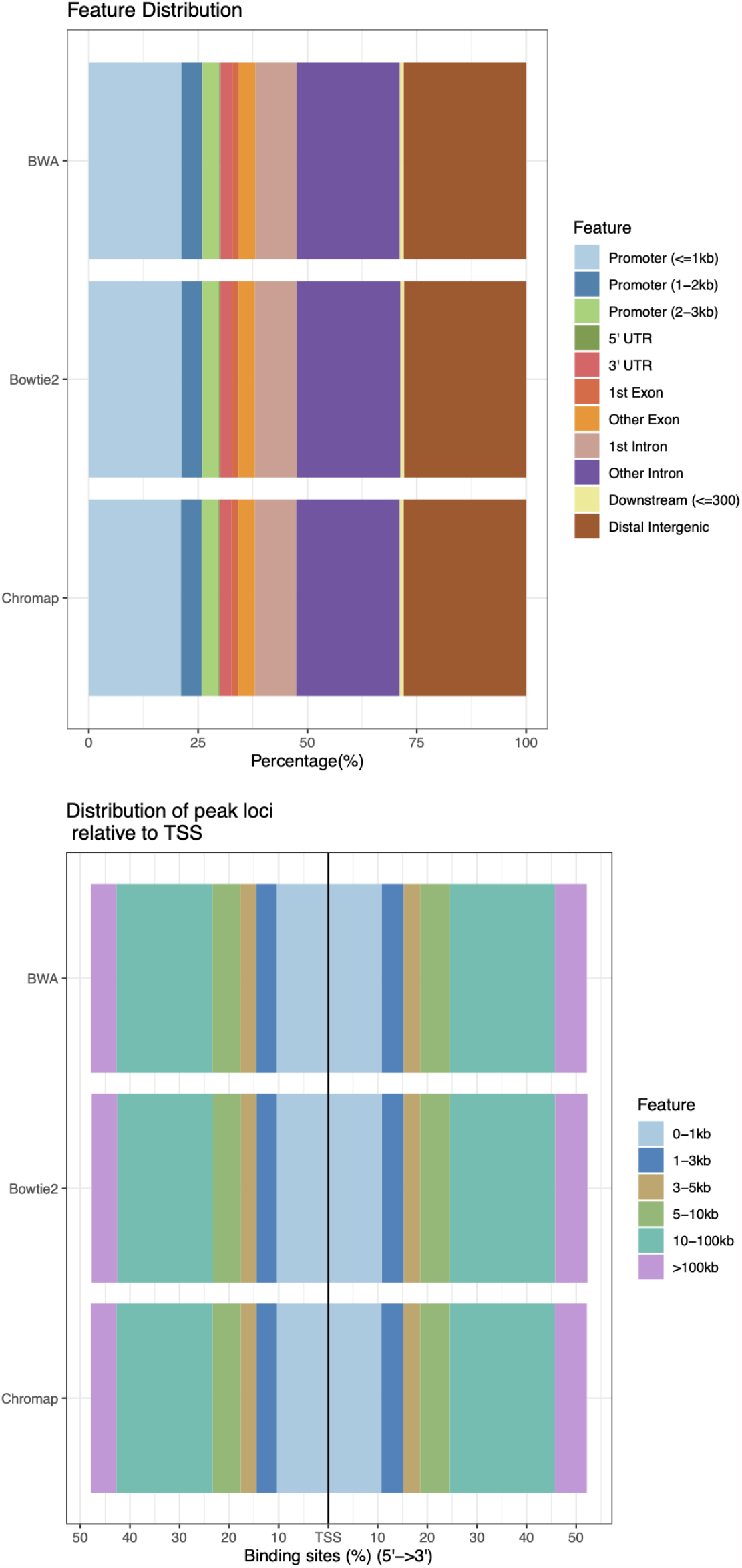
Annotations of peaks generated using MACS2 on BWA-MEM, Bowtie2 and Chromap alignments

**Supplementary Figure 2.**
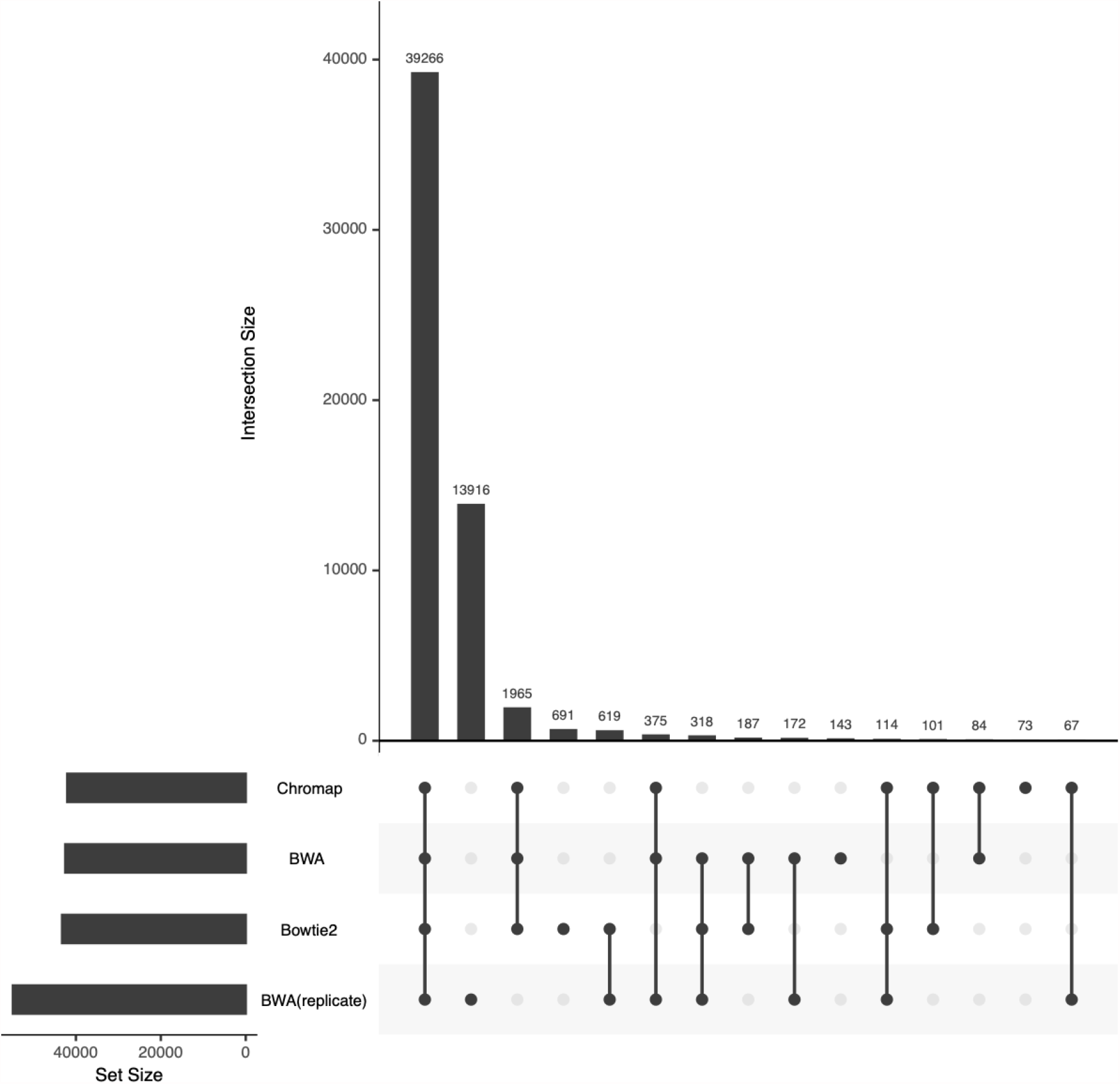
The number of overlapped peaks generated using MACS2 on BWA-MEM, Bowtie2, Chromap alignments of ChIP-seq replicate 1 and on BWA-MEM alignments of ChIP-seq replicate 2 (denoted as BWA (replicate))

**Supplementary Figure 3.**
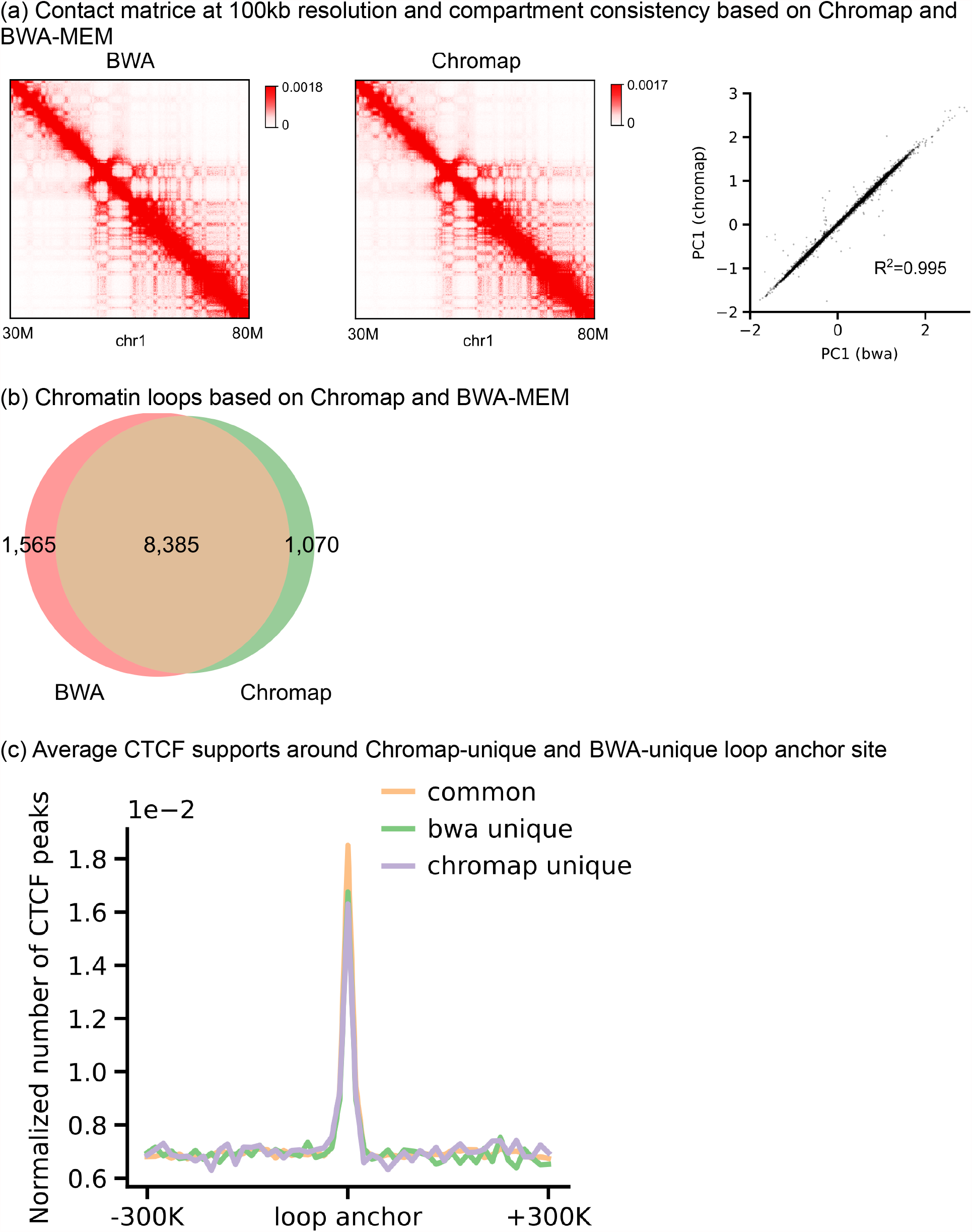
Evaluation of the Hi-C data.

